# Predictive coding during action observation - a depth-resolved intersubject functional correlation study at 7T

**DOI:** 10.1101/2021.08.30.458143

**Authors:** Leonardo Cerliani, Ritu Bhandari, Lorenzo De Angelis, Wietske van der Zwaag, Pierre-Louis Bazin, Valeria Gazzola, Christian Keysers

**Author notes:** contributed equally to the role of first author (Leonardo Cerliani, Ritu Bhandari, Lorenzo De Angelis). contributed equally to the role of last author (Valeria Gazzola, Christian Keysers).

## Abstract

While the brain regions involved in action observation are relatively well documented in humans and primates, how these regions communicate to help understand and predict actions remains poorly understood. Traditional views emphasized a feed-forward architecture in which visual features are organized into increasingly complex representations that feed onto motor programs in parietal and then premotor cortices where the matching of observed actions upon the observer’s own motor programs contributes to action understanding. Predictive coding models place less emphasis on feed-forward connections and propose that feed-back connections from premotor regions back to parietal and visual neurons represent predictions about upcoming actions that can supersede visual inputs when actions become predictable, with visual input then merely representing prediction errors. Here we leverage the notion that feed-back connections target specific cortical layers to help adjudicate across these views. Specifically, we test whether observing sequences of hand actions in their natural order, which permits participants to predict upcoming actions, triggers more feed-back input to parietal regions than seeing the same actions in a scrambled sequence that hinders making predictions. Using submillimeter fMRI acquisition at 7T, we find that watching predictable sequences triggers more action-related activity (as measured using intersubject functional correlation) in the parietal cortical area PFt at depths receiving feed-back connections (layers III and V/VI) than watching the exact same actions in scrambled and hence unpredictable sequence. In addition, functional connectivity analysis performed using intersubject functional connectivity confirms that these increased action-related signals in PFt could originate from ventral premotor region BA44. This data showcases the utility of intersubject functional correlation in combination with 7T MRI to explore the architecture of social cognition under more naturalistic conditions, and provides evidence for models that emphasize the importance of feed-back connections in action prediction.

## Introduction

The network of brain regions recruited by the observation of actions is relatively well described across humans and monkeys. In monkeys, using anatomical tracers, electrophysiology and fMRI, a temporo-parieto-frontal triad of mutually connected regions responds to the sight of hand actions (Nelissen et al. 2005, 2011; Rizzolatti and Sinigaglia 2016): regions lining the superior temporal sulcus (STS) are mutually connected with inferior parietal nodes (AIP/PFG) that are mutually connected with ventral premotor regions (F5a/c). In humans, exact homologies are difficult to establish, but studies and meta-analyses showed that observing goal-directed hand actions increases activity in lateral occipital/posterior temporal, parietal and premotor clusters that often peak in the lateral occipital cortex, parietal region PFt (similar to monkey PFG) and premotor region BA44 (similar to monkey F5a), and that these regions are also activated during the execution of similar actions (Gazzola and Keysers 2009; Caspers et al. 2010; Rizzolatti and Sinigaglia 2016).

While this network has long been implicated in participants’ ability to perceive the actions of others, a more recent hypothesis has been that they play a key role in *predicting* the actions of others. Reviewing recent neuromodulation studies confirms that perturbing activity in these regions interferes with the participants’ ability to predict upcoming actions in sequences (Keysers et al. 2018). Single cell recordings in parietal and premotor nodes in monkeys revealed that neurons responding to the sight of hand actions often have responses that differ based on the sequence of actions that can be predicted to follow (Bonini et al. 2010). Such recordings have also shown that the response to seeing a hand grasp behind an occluder increases when the animal can predict the presence of an object behind that occluder (Umiltà et al. 2001).

To map the brain regions encoding information about the sequence of motor acts that form complex action sequences, we video-recorded familiar sequences of acts that jointly form meaningful action sequences (Fig. 1a and Movie 1), and compared brain activity - measured using fMRI at 3T - between conditions in which participants viewed the motor acts either in their original (and thus predictable) sequence (Intact condition) or in a randomized (and thus unpredictable) order (Scrambled condition), (Thomas et al. 2018). We reasoned that nodes of the action observation network can either only contain information about individual motor acts or also additional information about the sequence in which acts organize into complex actions with more distal goals. If a node only represents individual motor acts, it should respond similarly whether these acts are in their natural order (Intact) or not (Scrambled). If it contains information at the sequence level, we would expect its activity to be sensitive to the order in which motor acts are presented, and thus respond differentially.

**Figure 1:**
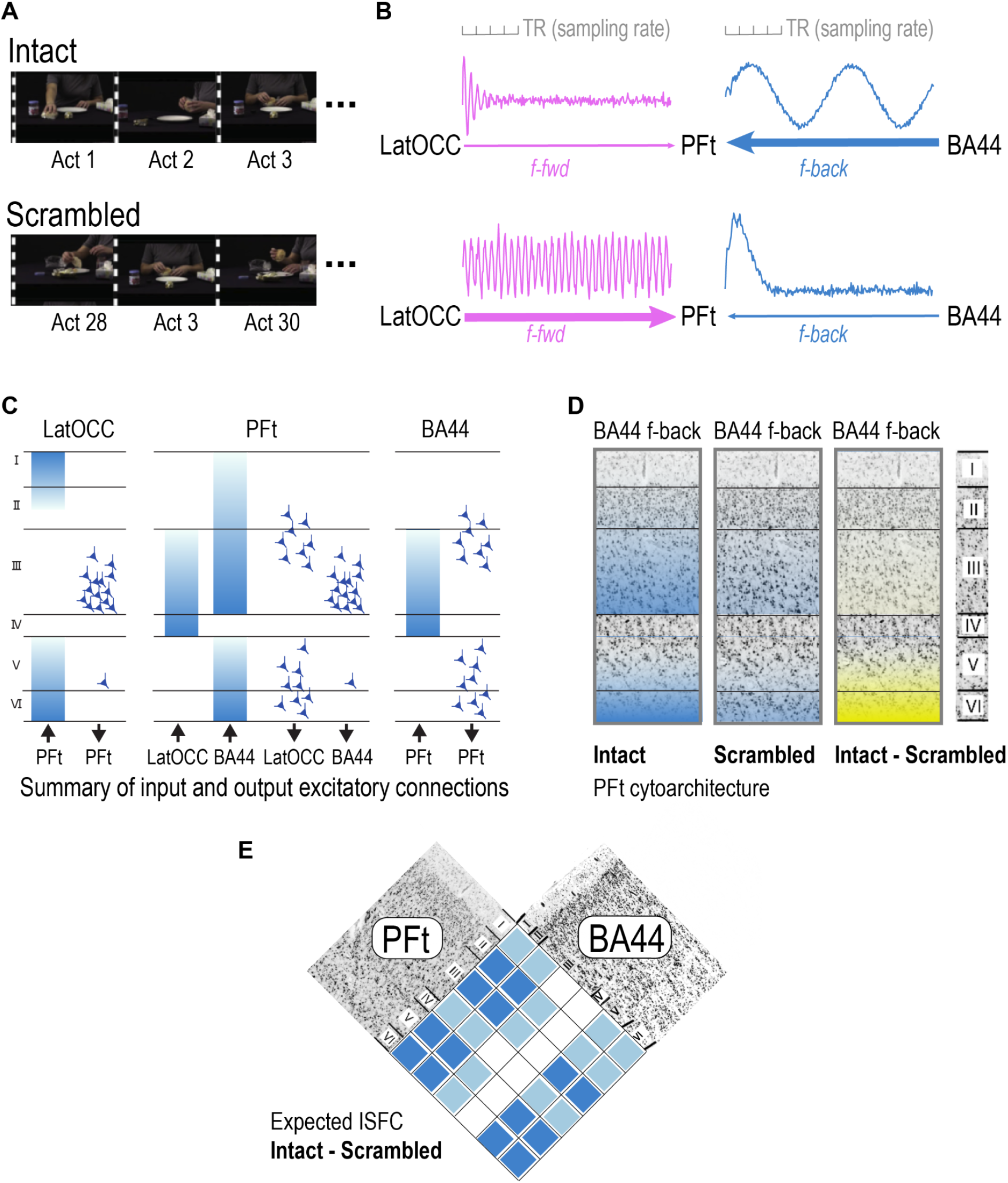
Stimuli and hypotheses. (a) We presented participants with movies of everyday hand actions lasting ∼1 min in length, filmed simultaneously with two cameras 45° apart, that were cut into ∼30 individual motor acts lasting ∼2 s (Table 1). In the intact condition, the motor acts were presented in their original order, but switching from one camera-view to the other at the transition between acts. In the scrambled condition, the acts were presented in randomized order. The change between camera-views were introduced in both conditions, because randomizing would otherwise have introduced visual transients in the Scrambled but not in the Intact movie. (b) Based on predictive coding, we hypothesize that **for Intact sequences** (top row), the parietal region PFt should receive comparatively little prediction errors from high level visual regions in the lateral occipital cortex (LatOcc). This is because in predictable sequences, only the first act cannot be predicted, and should thus trigger a strong prediction error, while subsequent acts become increasingly predictable, leading to waning prediction-error input to PFt. In contrast, PFt should receive strong and sustained predictive feedback from ventral premotor regions, and especially from BA44 (Keysers and Perrett 2004; Kilner and Frith 2008; Friston et al. 2011; Keysers and Gazzola 2014). Input from visual regions should be dominated by higher frequencies representing the individual motor acts presented every ∼2 s, while feedback from premotor regions would additionally include lower frequencies encoding sequence level information with periods up to the full ∼1 min of the movie (Honey et al. 2012; Thomas et al. 2018). **For Scrambled sequences**, PFt should receive sustained prediction errors from visual cortices while feed-back predictions from the premotor area should increasingly wane as participants realize that the sequence is unpredictable. (c) Adapting general organizational principles of the cortex summarized in Shipp (2007) by incorporating specific knowledge about the connectivity between ventral premotor and inferior parietal cortices obtained from monkey tracer studies (Gerbella et al. 2011) we can generate hypotheses about the layers in which the feed-forward information from LatOcc and the feed-back information from BA44 should arrive in PFt (gradients, middle panel), and where the cells in latOcc and BA44 from which this input originates are located (cells in the right and left panel, respectively). In addition, we can hypothesize where PFt signals back to LatOcc and forward to BA44 might terminate (gradients right and left panels), and where they might originate in PFt (cells in middle panel). Given that BOLD appears to be dominated by synaptic input rather than spiking output (Logothetis 2003), inputs (as illustrated by the gradients) would be expected to dominate the BOLD signals. Panel (c) is our own illustration of the schema in Shipp (2007) after modifying the gradients in PFt from BA44 by adding input to layer III based on Gerbella et al., 2011, and by aligning all layer thicknesses to those in PFt for readability although they vary across the brain regions. (d) Predictions about the depth profile of stimulus locked information (as measured using ISC) in PFt for Intact and Scrambled conditions based on (b) and (c). Intact conditions should be dominated by feed-back from BA44 triggering high ISC in layers III, V, VI (blue gradients in left column), while Scrambled conditions should show less feedback from BA44 (weaker blue gradients in the middle column). Accordingly, we expect the contrast between the ISC in the two conditions to be particularly clear in the deep layers (V/VI) and/or layer III with more ISC for the intact sequences (yellow gradient in the right column). This feedback information includes sequence level information unfolding over the tens of seconds of the entire movie (approximately between 0.1 and 0.015 Hz) and is thus slow enough to be well within the Nyquist limit of our fMRI sampling (TR=4.2 s or ∼0.24 Hz). In contrast, the feed-forward information representing prediction errors at each camera change (i.e. every 2 s on average), which should terminate in layer IV, and be stronger in the Scrambled condition, is too fast for its fluctuations to be accurately captures at our slower acquisition speed. Because intersubject functional connectivity depends on measuring the fluctuations over time of the signal rather than average BOLD activation, we hypothesize this difference in layer IV to be difficult to measure using the methods we employ(Nastase et al. 2019).. (e) Predictions about the different (Intact vs Scrambled) functional connectivity between PFt and BA44 and LatOcc - measured using ISFC - based on the hypothesis described in (b) and (c): we expect PFt to show increased connectivity with BA44 for Intact > Scrambled with a feedback pattern of connectivity, while we expect it to show increased connectivity with LatOcc for Scrambled > Intact in accordance with a feedforward pattern of connectivity. The specific stratification of the feedback depends on whether the feedback would originate from BA44 cells in layers II/III or layers V/VI, and whether it would terminate more around layer III or layers V/VI, all of which would be anatomically plausible (Shipp 2007; Gerbella et al. 2011).

To identify nodes with BOLD signals containing sequence level information, we used intersubject correlation (ISC, Nastase et al. 2019) to map and quantify stimulus locked information in the brain. Lerner et al. (Lerner et al. 2011) had pioneered this approach to identify sequence level information in language processing by having participants listen to narrated stories either in their intact order, or in scrambled order, in which for instance the same words appeared in random order. They found that while auditory cortices show similar levels of synchronization across listeners whether the story was heard intact or scrambled, frontal brain regions synchronized across listeners much more in the intact condition, showing that frontal regions integrate information across words into meaningful sequences. Adapting this elegant approach to actions, we found that left PFt, and to a lesser extent BA44, showed higher ISC for intact than scrambled hand action sequences, establishing that these regions contain sequence level information (Thomas et al. (2018)). Temporal and occipital visual regions showed high ISC for Intact and Scrambled sequences, and thus encoded information about the observed acts, but did not show higher ISC in the intact compared to scrambled condition, suggesting that they do not contain significant levels of sequence level information. Together, these results sketch a hierarchical picture in which the temporal/occipital lobes process observed actions, but at the relatively short temporal scales of single motor acts, while PFt and BA44 process these acts in ways that integrate that information into longer - and arguably more meaningful - sequences.

How this network integrates individual motor acts into meaningful sequences is in the focus of a number of theoretical papers (Keysers and Perrett 2004; Kilner and Frith 2008; Friston et al. 2011; Keysers and Gazzola 2014), but empirical data to support these notions remains scarce. Early work emphasized feed-forward information flow in a hierarchical structure from the temporal lobe to parietal and premotor regions where action understanding and action prediction then occur by triggering premotor action plans in mirror neurons. For instance, Gallese et al. in their seminal 1996 paper wrote: “The superior temporal sulcus representation would provide, in this case, an initial description of hand object interactions that would then be sent to F5 and matched with the ‘motor vocabulary’ of that area. […] When an individual is carrying out a goal-directed action, he ‘knows’ (predicts) its consequences. [Premotor] Mirror neurons could be the means by which this type of knowledge can be extended to actions performed by others” (Gallese et al. 1996, p606). Indeed, MEG data supported this notion by measuring increasing latencies across these nodes (Nishitani and Hari 2000). A problem with this feed-forward view is that it implies that for a person to respond to the actions of others, the sight and sound of that action would need to trigger activity in a feed-forward stream through high level sensory, parietal and then premotor regions. This would imply delays of ∼200ms between witnessing an action and executing a response. However, pianists have been shown to adjust their own key presses to within 30ms of those of others (Keller et al. 2007) - an order of magnitude too fast for such a simple feed-forward architecture (Keysers and Gazzola 2014). More recently, it has been argued that the action observation system could instead use Bayesian inference in a predictive coding system that combines the feed-forward transmission of prediction errors with feed-back transmission of predictions across the mutually connected nodes of the system (Keysers and Perrett 2004; Kilner and Frith 2008; Friston et al. 2011; Keysers and Gazzola 2014). Specifically, we proposed that through their mutual connectivity, the premotor-parietal system, which orchestrates the temporal unfolding of complex action sequences during motor control, has the sequence level information spanning tens of seconds that psychologists call ‘motor schemata’, while the occipito/temporal nodes only represent individual actions, and receive predictions from the parietal lobe, and send prediction errors to the parietal lobe (Thomas et al. 2018). If the predictions match the incoming visual information (i.e. small or no prediction error) the sensory input from visual areas is attenuated - an attenuation we indeed measured in ERP responses over the occipital lobe (Thomas et al. 2018). If they do not match, the difference (large prediction error) is sent forward to parietal and premotor regions where they can activate alternative motor schemata (i.e. action sequences) that better match the visual input of the observed action sequences. If observed acts continuously violate expectations, participants report stopping to generate predictions and premotor cortices may then stop to send such predictions backwards towards parietal cortices (Thomas et al. 2018). A core - and testable - tenet of this predictive coding hypothesis is that the sight of actions organized in predictable sequences should induce more feed-back information flow instantiating predictions from BA44 to PFt while scrambled sequences should induce more feed-forward information flow instantiating prediction errors (Fig. 1b). The traditional, hierarchical feed-forward view of action observation would not predict such an increase of feed-back information as subsequent motor acts in the sequence become predictable. In terms of time scales of information, the feed-forward input from visual regions would contain prediction errors at each transition across acts, in the scrambled condition in particular, that would have relatively fast frequencies (every ∼2 s), while the feed-back input from premotor regions would contain a mix of frequencies that span from signals that predict the upcoming act (every ∼2 s) and sequence level information at lower frequencies (with periods T up to ∼1 min). Such a mix of frequencies has been observed in similar experiments on language (Honey et al. 2012).

To summarize these predictions from the point of view of PFt, we thus have the following expectations (Fig. 1b): because participants do not know what sequence will be presented, PFt should initially receive visual input about the first acts of a sequence, an input that then rapidly wanes as PFt increasingly suppresses expected visual input. This input is dominated by the high frequency signals (T≈2s) of the individual motor acts. PFt should also receive predictions from BA44 in a broader frequency range, that include short term predictions about the unfolding of the current act (T≈2 s) to long term predictions about the entire sequence (T≈1 min). These BA44 feed-back predictions should remain high throughout the sequence, and dominate the overall signal during the Intact sequences. In contrast, for the Scrambled sequences, predictions cannot suppress the randomized presentation of the acts and PFt should thus receive sustained feed-forward prediction error input from visual regions at each camera change. Predictions from BA44 are likely to subside as participants realize that the sequence is not predictable, leading BA44 feedback to wane, so that the BOLD signal overall should be dominated by the feed-forward signal. Because the BOLD signal appears dominated by synaptic input over spiking output (Logothetis 2003), we neglect what signals PFt is likely to output to other regions in these considerations. Considering what we know about the laminar distribution of feed-forward and feed-back inputs from a variety of sensory brain regions organized in hierarchical fashion (Fig 1c, adapted from(Shipp 2007)), feed-forward input from visual regions should terminate in middle layers of PFt (layer IV) and feed-back input from BA44 should terminate mainly in deep layers of PFt (layers V/VI). However, monkey tracer studies have shown that the connections from F5a (the likely monkey homologue of BA44) to parietal regions similar to human PFt terminate not only in deep layers V/VI but saliently also to layer III (Gerbella et al. 2011). Such deviation from the prototypical feedback connectivity pattern described by Felleman and Van Essen in the visual system (Felleman and Van Essen 1991) is indeed not usual for fronto-parietal connectivity across association areas (Rozzi et al. 2006; Gerbella et al. 2010, 2011). We can thus reformulate these predictions about the direction of information flow as predictions about the depth at which stimulus-locked information should dominate the BOLD signal in the Intact and Scrambled conditions if our model of predictive coding is true (Fig. 1d). With regard to feed-back information flow, which should be dominant in PFt in deep layers (layer V/VI) and/or in layer III, we expect more for Intact than Scrambled conditions, so that the contrast Intact-Scrambled should show increased ISC in layers III, V and VI. This feed-back information flow contains information at the sequence level, that fluctuates over tens of seconds, and our relatively slow 7T acquisition (TR=4.1s, Nyquist limit T=8.2s) should thus be adequate to capture its fluctuations over time, which is the basis for intersubject correlation analyses(Nastase et al. 2019). In contrast, the feed-forward information flow, that caries prediction errors, would peak at every camera change (T≈2s) in the Scrambled condition, and fluctuate faster than the 7T fMRI acquisitions we will use in this study (TR=4.1s, Nyquist limit T=8.2s). There might thus be higher *average* BOLD activity over the duration of the movies in the Scrambled condition in layer IV, where feed-forward should terminate, but the ISC method we use does not investigate average BOLD activity, but the synchronization of activity fluctuations between participants(Nastase et al. 2019). However, we expect to fail to measure the second by second fluctuations of such prediction errors accurately enough for ISC to evidence increased synchronization in layer IV for the Scrambled condition. Accordingly, for ISC, the most robust observable effect derived from the difference across conditions should be the increased stimulus-related information in deep layers (V/VI) and/or layer III for the Intact sequence. Such an increase is not predicted by traditional models of action observation that do not emphasize the role of feed-back connections, and can thus help adjudicate between these views.

Pioneering studies in the early visual cortex have shown that acquiring fMRI at submillimeter resolution at high-field strength (≥7T) can indeed test predictive coding hypotheses (Kok et al. 2016; Lawrence et al. 2018, 2019a, b; Finn et al. 2020; Aitken et al. 2020). Recent work in particular confirms that activations in deep layers is indeed measurable when feedback predictions about upcoming sensory events are likely to occur in the visual system (Kok et al. 2016; de Lange et al. 2018; Aitken et al. 2020), while activation in superficial layers seem to be measured when there is a mismatch between predicted and observed input and attentional signals (de Lange et al. 2018). Unfortunately, experiments testing whether deep layers carry signals that could be related to predictions have so far been restricted to early visual cortices and we do not know whether the same might be observed in parietal and premotor regions (Finn et al. 2020). The additional challenge of BOLD signal integration across layers due to veins increases the complexity of the question (Markuerkiaga et al. 2016; Huber et al. 2017). Here we acquired brain activity at 7T with an isotropic resolution of 0.8 mm in 9 participants, planning a partial-volume acquisition to include PFt, BA44 and LatOcc cortex of the left hemisphere (Supplementary Figure 1), while they observed intact and scrambled actions to test the validity of predictive coding models of action observation, specifically by testing for increased stimulus-dependent information in deep layers of PFt when seeing intact vs scrambled sequences. We choose PFt as our main region of interest because it is at the crossroad between feed-forward input representing prediction errors and feed-back information from BA44, and because it has a layering that is somewhat similar to the better explored visual system, while BA44 and its dysgranular organization (Amunts et al. 1999; Amunts and Zilles 2012) would make the same approach more tentative.

Methodologically, as in our previous 3T work (Thomas et al. 2018), we use intersubject correlation (ISC) to localize sequence-level information, and intersubject functional correlation (ISFC) to quantify information sharing across nodes in this study for two main reasons. First, to generate sequence level brain activity, we need longer sequences of actions (e.g. preparing breakfast) in the order of ∼1min. Such stimuli are too long for traditional block-design fMRI experiments, but are ideal for ISC analyses (Nastase et al. 2019). Second, quantifying functional connectivity between brain regions within subjects is complicated by the fact that a sizable proportion of the BOLD signals are formed by noise that is often shared across brain regions (e.g. respiration and motion (Liu 2016)). That two brain regions synchronize their BOLD signal can thus be due to true exchange of stimulus relevant information across those regions or to the presence of common noise. Intersubject functional correlation (ISFC) provides a way to circumvent the connectivity-measure-inflating effect of such common noise across brain regions (Simony et al. 2016; Nastase et al. 2019) by leveraging the fact that BOLD signals in each voxel of the brain can be seen as a combination of (i) stimulus locked signals that represent stimulus-relevant information and (ii) signals unrelated to the stimulus (e.g. daydreaming and noise) that will not be stimulus locked. By correlating the signal in region PFt of one participant with the average time course of all other participants in region BA44 or LatOcc, ISFC can then isolate stimulus-locked information shared across these regions, because noise or stimulus unrelated thoughts would be unlikely to occur systematically at the same time in different participants, and therefore would average out and not correlate in time across participants. Here we will thus leverage ISC to test the presence of more ISC in layers III or V/VI of PFt for intact over scrambled sequences and ISFC to test that stimulus related information in PFt is more shared with BA44 during intact vs scrambled sequences. Finally we can also leverage ISFC to explore whether information at depths compatible with Layer IV of PFt may indeed be more shared with LatOcc regions during scrambled sequences, although we hypothesize that this analysis will be limited by our ability to detect fast stimulus-dependent fluctuation of the BOLD signal in LatOcc due to our fMRI sampling frequency.

## Methods

### Participants

Data were acquired from 14 subjects (10 males, 4 females, Age mean∓std = 24.9∓2.8, range=21-30). Three subjects were excluded due to missing data and two additional subjects were removed due to excessive head motion. Therefore 9 subjects (6 males, 3 females, Age mean∓std= 24.8∓2.6, range=21-30) were included in our analyses. Upon arrival the subjects were screened for MRI compatibility and were familiarized with the MRI setup. All subjects reported being right handed and having no psychological or neurological disorders. All of them had a normal or corrected to normal vision. Subjects were compensated 10 Euros/hour. The study was approved by the Lab Ethic Review Board of University of Amsterdam (2016-BC-6837) and all subjects signed the informed consent before the beginning of the experimental session.

### Stimuli

Eighteen movies containing different daily actions (e.g. preparing sandwiches with butter and jam; see Table 1 for the full list) were simultaneously recorded by two video cameras (Sony MC50, 29 frames/s) at an angle of 45 degrees. The videos were edited using ADOBE Premiere ProCS5 running on Windows. Each movie was subdivided into shots containing one meaningful motor act each (e.g. taking bread, opening the butter dish, scooping butter with a knife, etc.). This was done on recordings from both camera angles. These motor acts (mean/standard deviation duration 2s ± 1s) were then assembled to build two types of ∼1minute long stimuli (Fig. 1).

**Table 1:**
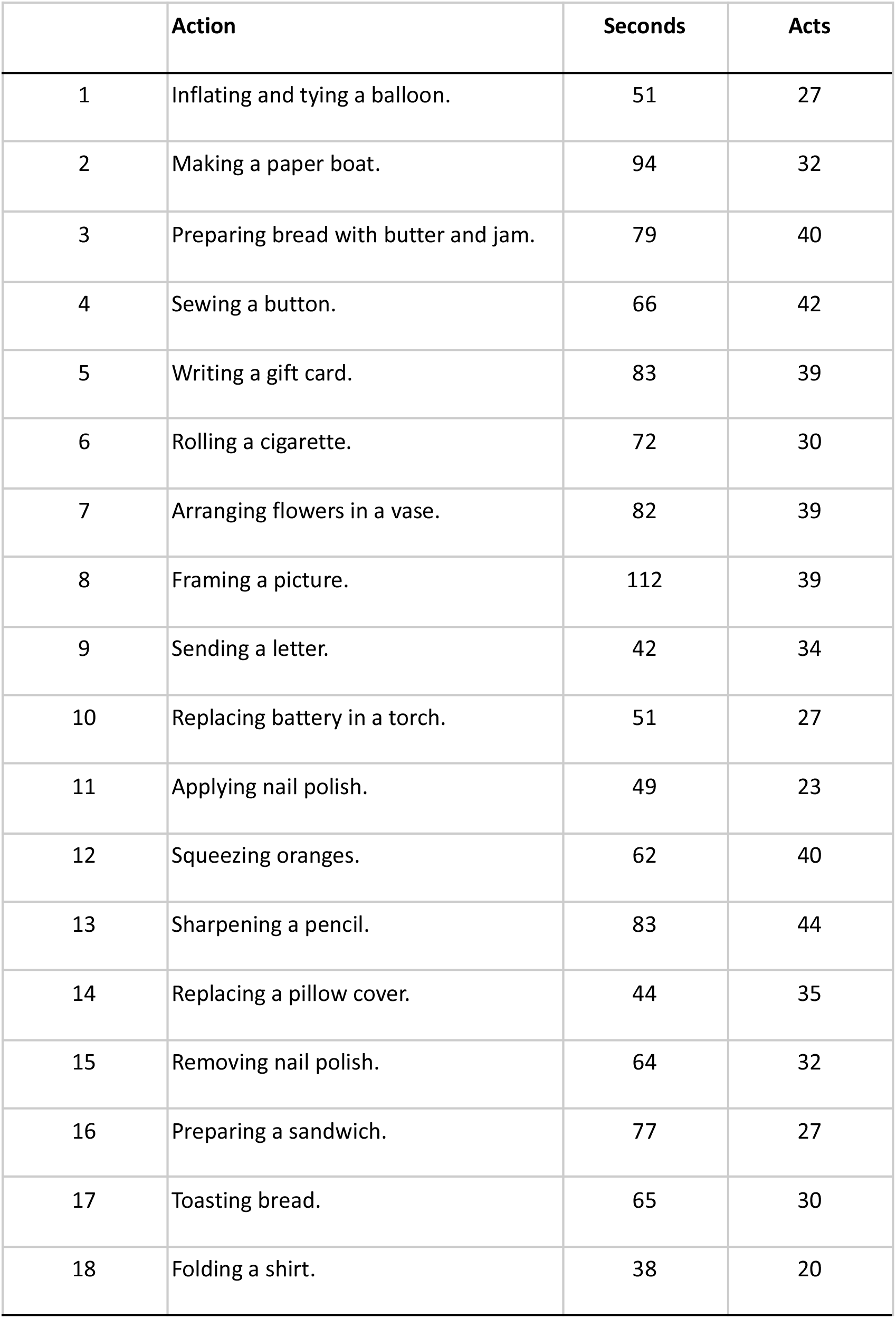
List of sequences used as stimuli with total duration in seconds and number of motor acts.

For the *Intact* (I) presentation, the natural temporal sequence in which the acts were recorded was maintained, but a camera angle change was introduced between every two consecutive acts by alternate sampling from the recordings of the two cameras. In the *Scrambled* (S) versions, the acts remained the same, but the order of the acts was randomly re-arranged, and a camera angle change was introduced between every two consecutive acts. Camera angle changes were imposed at each act transition in both types of movies to compensate for the visual transients that would otherwise be present only in the scrambled movies.

This resulted in 18 intact and 18 scrambled movies. These movies were presented to the subjects in 4 blocks. Block 1 (220 fMRI scans), 2 (160 fMRI scans) and 3 (190 fMRI scans) each consisted of 5 unique intact movies and the corresponding 5 scrambled movies. Block 4 (145 fMRI scans) consisted of the remaining 3 intact movies and the 3 corresponding scrambled movies. The stimuli in each block were separated by inter-movie interval varying from 8 to 12 s. The unique combination of videos in each block was kept constant across participants to ensure that each subject always viewed the scrambled version of a movie before its intact version.

### Experimental procedure

Each subject was invited twice, around one week apart (Mean=5 days, Range = 2 -10 days). On the first day, they viewed blocks 1 and 2 of Intact and Scrambled movies, in two separate but consecutive 3DEPI fMRI data acquisitions. This was followed by the acquisition of anatomical images (MP2RAGE and T1-3DEPI). Subjects then viewed the same 2 blocks of the movies again while we collected new 3DEPI images. The order of the blocks was pseudo-randomized across subjects. During the second day, subjects saw blocks 3 and 4 of Intact movies and their temporally Scrambled versions. This was followed by a T1-3DEPI image acquisition followed by repetition of the blocks. Similar to the first session, blocks were randomized across subjects. Videos were presented using the Presentation software (Neurobehavioral Systems, Inc., Albany, CA, USA). No behavioural response was required during the four sessions, but participants were instructed to carefully observe the videos. We ensured that subjects were not in the scanner for more than 90 minutes during each session.

### Image acquisition

All MRI images were acquired on a Philips 7T Achieva at the Spinoza Center in Amsterdam (https://www.spinozacentre.nl/) using a 32-channel receive and 2-channel transmit volume coil (Nova Medical, USA). On each experimental day, we acquired 4 runs of partial fMRI brain images, and one anatomical (T_1_23DEPI) image, with the same field of view and in-plane resolution of the 3D EPI fMRI images. Images were centered on the inferior posterior parietal lobe (rectangle in Figure 2A), but also covered lateral portions of the temporal, occipital and inferior frontal lobe. We manually adjusted the bounding box to approximately cover the same brain regions in all subjects. On the first day only, we additionally acquired one high-resolution whole-brain anatomical image (MP2RAGE). All images were acquired in sagittal orientation.

**Fig 2.**
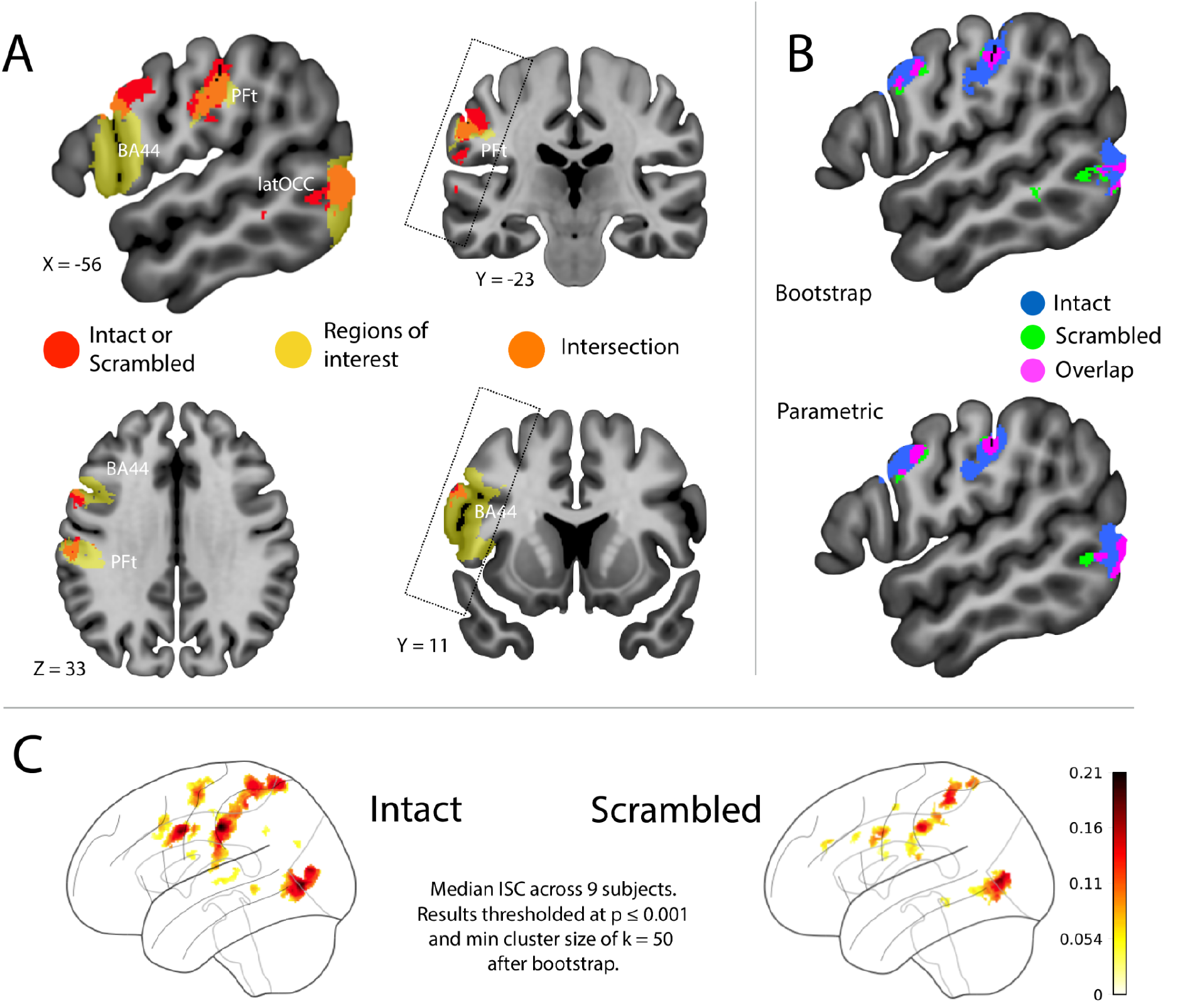
Results of the Localizer-ISC. The first ISC was carried out in MNI space, to define patches of interest with brain activity relating to hand action observation during either Intact or Scrambled movies, to be later used in the second, depth-resolved ISC in native space. **A**. Results (significant voxels) for a logical OR between the contrasts ISC_Intact_ > 0 *or* ISC_Scrambled_> 0 (red and orange), overlaid onto the initial regions of interest (yellow): BA44 and PFt from the Anatomy toolbox, lateral occipital (latOCC) from the Harvard-Oxford cortical atlas. The latOCC was used as an anatomical region that corresponds to the posterior mid temporal gyrus clusters often observed in previous studies of action observation. Significantly active voxels co-located with voxels in the regions of interest are denoted in orange. The rectangle crossing the coronal images approximates the field of view common to our participants afforded by our scans. Lack of ISC outside of this box should and cannot be interpreted as suggesting that these regions do not show significant ISC. The exact extent of the field of view across subjects can be found in Fig S1 **B**. Comparison of the results obtained using bootstrap (*p*_unc_<0.001, k=50 minimum voxels per cluster) and parametric inference (*p*_unc_<0.001, k=50 minimum voxels per cluster). Scrambled activity was overall associated with smaller clusters mostly encompassed in regions activated by Intact movies. **C**. Median ISC parameter estimates across 9 subjects. Significance was established according to the null distribution of median ISC values obtained using bootstrap. Median ISC values lying beyond the 99.99th percentile of the null distribution (i.e. p ≤ 0.001) were considered significant. In addition, a minimum cluster size of k=50 voxel was applied to determine significance.

#### fMRI acquisition

3D EPI sequence at 0.8 × 0.8 × 0.8 mm resolution, FOV 169 × 150, 30 slices, reconstructed at 0.75 mm, each 3D volume acquired in 4.11 s, EPI factor = 27, SENSE(AP) = 3.9, flip angle = 18º, TR/TE = 54/28 ms.

#### partial FOV anatomical acquisition

T_1_23DEPI (van der Zwaag et al. 2018a, b) sequence at 0.8 × 0.8 × 0.8 mm resolution, FOV 169 × 150, 30 slices, reconstructed at 0.75 mm, SENSE(AP) = 3.9, inversion time (inv1/inv2) 0.8/2.7 s, flip angle 20/16º, TR/TE = 52/25 ms.

#### whole-brain anatomical acquisition

MP2RAGE (Marques et al. 2010) sequence at 0.64 × 0.64 × 0.64 mm resolution, FOV 220 × 220, 256 slices, reconstructed at 0.625 mm, inversion time (inv1/inv2) 0.8/2.7 s, flip angle 7/2º, TR/TE = 6.2/2.2 ms.

### Image preprocessing

In our preprocessing pipeline, we aimed at preserving the ability to detect the cortical depth of the BOLD signal by minimizing the amount of interpolation steps for the fMRI images, and planned the other necessary preprocessing steps involving anatomical images around this aim. Specifically, the native space of the fMRI images was kept as the reference for all subject-level and group-level inter-modal registration and analysis, an approach also adopted by previous studies (Huber et al. 2017) to preserve the layer-specific information in high-resolution fMRI images.

The whole preprocessing pipeline can be accessed at the official github repository for this study (https://github.com/ldeangelisphys/layerfMRI). The scripts (in bash, Python, R) employ routines from a variety of MRI image processing packages: FSL (https://fsl.fMRIb.ox.ac.uk/fsl), nighres (https://nighres.readthedocs.io/) ANTs (https://stnava.github.io/ANTs/), ITK-SNAP and associated command-line utilities (http://www.itksnap.org/).

#### Image reconstruction

The PAR/REC stacks exported from the Philips scanner were converted to NIFTI images using dcm2niix v1.0.20201102 (https://github.com/rordenlab/dcm2niix) and reoriented to the standard RPI coordinate system using fslorient2std, to enable inspecting the images in a familiar orientation. Facial information was removed from whole-brain anatomical images using pydeface (https://github.com/poldracklab/pydeface). Finally, the T1w and the quantitative T1 map images were reconstructed from the two inversions of the MP2RAGE acquisitions using the nighres `nighres.intensity.mp2rage_t1_mapping` function and the appropriate acquisition parameters. All the other parameters of the functions were left at the default values. The T1 images were subsequently used to estimate the cortical depth and the registration with the fMRI images.

#### Skull stripping

Removal of the skull was necessary to carry out image registration across MRI modalities within subject and with the MNI template. A precise extraction of the cortical sheet was instead carried out later (see below “Cortical depth estimation”). For the full anatomical acquisition, we obtained the best results by first correcting the second inversion image of the MP2RAGE (inv2) for bias field using ANTs’ `N4BiasFieldCorrection`, then feeding this and the quantitative T1 map into the nighres’ `nighres.brain.intensity_based_skullstripping` function to carry out skull stripping. In the partial brain anatomical images, we carried out the bias field correction on the mean inv2 acquisition for each participant, and then thresholded this image to an intensity of 1.5e5. In the partial brain fMRI images, we thresholded the intensity-normalized mean fMRI image to either 0.05 or 0.1 (2/9 and 7/9 participants, respectively), and then generated the brain mask using `ants.get_mask` with `cleanup` option set to 1 (erosion with a radius of 2 voxels). These values of intensity threshold were manually chosen based on visual inspection of the skull stripping, after observing that the default parameters would produce a suboptimal result (i.e. either removing brain tissue or not identifying voxels in the skull for removal) which would have had consequences during image registration.

#### FMRI image preprocessing

FMRI images underwent a minimal preprocessing pipeline, which consisted of brain extraction, motion correction to the middle image of each run using FSL MCFLIRT and detrending using a gaussian-weighted line as implemented in FSL FEAT. To preserve the cortical depth specificity of the fMRI signal, we did not apply spatial smoothing to the images that were used for the main analysis related to the cortical depth. Instead we applied a 6mm FWHM spatial smoothing only for the initial group-level analysis which served to identify the regions of interest later used in the cortical depth analysis. Details are provided in the Analysis section.

#### Registration

All registrations between anatomical (T1w) images were carried out using ANTs’ SyN algorithm (symmetric diffeomorphic registration preceded by an affine transformation) with mutual information as optimization metric (Avants et al. 2008, 2011; Klein et al. 2009) as implemented in ANTsPy. For the registration between fMRI and partial FOV anatomical image, we used the provided version of the SyN algorithm optimized for BOLD images (SyNBoldAff). The registration between the partial anatomical image (T1w) and the fMRI image was estimated using the mean fMRI image of each run and the partial anatomical image acquired in the same session. Additionally to these within-subject registrations, we also estimated the registration between the full anatomical image and the standard MNI (the 1 mm isotropic `icbm152_2009_brain.nii.gz` provided in nilearn - https://nilearn.github.io/) using the same SyN algorithm and optimization metric.

#### Cortical depth estimation

The estimation of the cortical depth from the high-resolution whole-brain anatomical images was carried out with the functions provided by nighres. This complex procedure involves several steps which have been extensively detailed in previous works (Bazin et al. 2014; Huntenburg et al. 2018). Here we briefly describe the pipeline adopted for our images.

The skull stripped version of the inv2 and quantitative T1 map were obtained using the brain mask from the previous skull stripping of the bias field corrected inv2 image. We then estimated two probability maps: one for the dura mater (based on the inv2 image), one for the sulcal ridges (based on the T1 map). Together with the initial brain mask, these additional maps provided further information for carrying out brain segmentation (Bazin et al. 2018) of the whole-brain anatomical images.

Brain segmentation was then carried out using a Multi-compartment Geometric Deformable Model (MGDM) (Fan et al. 2008; Bogovic et al. 2013) implemented in the `nighres.brain.mgdm_segmentation` function. The inputs to the function were the skull-stripped T1 map and a stack containing the dura and sulcal probabilistic maps. Since this procedure is based on labelled atlas priors (provided in nighres), the GM output map refers only to the cortex and excludes deep brain nuclei. The boundaries of the cortex are then extracted based on the results of the segmentation using a single object Geometric Deformable Model, and the cortical ribbon is reconstructed using the CRUISE algorithm (Han et al. 2004).

The CRUISE reconstruction results in the definition of the boundaries between GM/WM and GM/CSF, which are then used to estimate cortical depth using a volume-preserving method (Waehnert et al. 2014, 2016; Trampel et al. 2019). Note that per the defaults of nighres (see https://nighres.readthedocs.io/en/latest/laminar/volumetric_layering.html), cortical depth values - ranging from 0 to 1 - are higher towards the pial surface. We maintain this convention in presenting the results and in numbering the bins which will be used to pool voxels within different cortical depth ranges.

### Analytic strategy

We conducted our fMRI analysis in the framework of inter-subject correlation (ISC - Hasson et al. 2004) and the related inter-subject functional connectivity (ISFC) (Simony et al. 2016; Nastase et al. 2019). ISC is a relatively novel analytical paradigm where the significance of a brain region for a certain task is determined by estimating the similarity - using Pearson correlation - of the brain activity between different participants involved in the same task over time - e.g. while watching the same movie - rather than by regressing a hemodynamic model of the task on brain activity.

Generally, ISC is carried out at the voxel level after alignment of the entire 4D fMRI volume in a common space like the MNI. This however represented a limitation for the setting of our experiment, since the spatial interpolations required to move the native fMRI data into a common space (coregistration) decrease the spatial resolution of the BOLD signal. In order to benefit from the ISC analytic strategy in our data, while at the same time retaining the potential to resolve the fMRI signal as a function of cortical depth, we devised a procedure that we denominate “dual-ISC”, which allows an ISC analysis to be carried out on the fMRI data in native space.

The **first ISC** is carried out in MNI space after smoothing the fMRI data. The results from this ISC are used to identify the locations on the cortex where brain activity is significantly consistent across subjects during the observation of either Intact or Scrambled movies. The group-level binary masks of the clusters identified in this analysis are then transformed into the single subject native space, where they are used to estimate an average time course for each cluster using the original, non-smoothed data. Crucially, within each cluster the time course averaging is carried out separately for voxels at different cortical depths. This allows to carry out a **second ISC**, where the unit of analysis is not each single voxel’s fMRI time course, but rather the averaged time course for all the voxels within a certain cortical depth bin in each of the clusters of activity. Similar to spatial smoothing in conventional-resolution fMRI, the averaging also provides a way of denoising the fMRI signal before conducting the analyses. We will now describe the steps of this procedure in detail.

### First ISC - Localizer

The preprocessed fMRI images of each run for each subject were spatially smoothed with a gaussian kernel of 6 mm FWHM (8 times the original voxel size). The previously estimated transformations (fMRI to partial T1w, partial to full T1w, full T1w to MNI) were aggregated into one single transformation using `ants.apply_transforms`, to limit the amount of interpolation steps. This composite transformation was applied to the 4D images using linear interpolation. This allowed us to carry out the first standard voxelwise ISC analysis to define a group-level localizer of brain activity associated with the observation of intact or scrambled movies, without biasing the analysis for one of the others.

We then used the registered 4D images to perform an InterSubject Correlation (ISC) analysis. This approach requires correlating the time courses recorded in every voxel across different subjects to identify those that are synchronized across subjects, therefore reflecting shared stimulus-dependent fluctuations across subjects (Nastase et al. 2019). We use ISC to determine which regions are involved in the processing of either intact or scrambled clips that we show to each participant.

The ordering and timing at which the clips are shown was randomized across participants. Before carrying the intersubject correlation, we then extract the segment (i.e. series of volumes) corresponding to each clip, we standardized each segment (removed mean and divided by the standard deviation) and concatenate them following the same order for every participant. This generated one 4D dataset containing all the data segments corresponding to intact movies concatenated together in a cardinal order and one containing all the scrambled movies. Note that although for both conditions each movie was displayed twice, such repetitions are treated as two independent movies and concatenated accordingly.

Following this processing of the data, we ran ISC in a leave-one-out approach. This procedure consists in correlating the time course of a voxel for one subject with the average time course corresponding to all the other subjects (Nastase et al. 2019). We then use a non parametric approach (bootstrap) to identify the brain regions with a significant ISC (Chen et al. 2016). Given the relatively low sample size (9 subjects), we also perform a traditional parametric t-test, which is predicted to produce less false positives than non-parametric metrics in the case of limited sample sizes (De Angelis et al. 2021). Having observed that both thresholding methods yield consistent results, we threshold the ISC map with the bootstrap method, with a p-threshold of α = 0.001 and a minimum cluster size of k = 50.

Having observed that both thresholding methods yield consistent results, we estimated the significance of median ISC values using bootstrap (Chen et al. 2016) across 5000 subject-wise resampling (with replacement). ISC values were considered significant if they lied outside the mid 99.95% (equivalent to a two-tailed p ≤ 0.001) of the bootstrap distribution, after adjusting (i.e. shifting) the distribution for the median ISC of the actual sample. We then apply the cluster size filter with a custom made python code based on the sklearn-image library (cluster connectivity = 2, as in SPM). This result was used to produce a binary mask in MNI space encompassing the brain locations where a significant degree of synchronized activity was found across participants during the observation of either intact or scrambled movies representing goal-directed actions.

### Second ISC - Cortical depth specific

The localizer in MNI space was taken into the native fMRI space of each run for each subject using the inverse of the previously estimated composite transformation from fMRI to MNI space. Concerning this and all other transformation to the native space, since we were transforming volumes containing labels (either significantly active regions from ISC1, atlas regions or cortical depth) the information of which should be preserved as such (i.e. integers), we used a nearest neighbour interpolation (`genericLabel` option in ANTsPy, recommended by ANTs team for images of this kind).

As detailed in the Introduction, we had a strong anatomical hypothesis about the location of the regions where we expected to observe a cortical-depth specific effect of observing intact vs. scrambled goal-directed actions. Specifically, we focused our investigation of the inferior posterior parietal lobe on area PFt (Caspers et al. 2006), of the ventral premotor cortex to the posterior inferior frontal gyrus Brodmann area 44 (Amunts et al. 1999) and the higher-order lateral visual cortices. We defined our regions of interest using a logical AND between the functional localizer obtained by the first ISC and the cytoarchitectonic maps from Juelich (Eickhoff et al. 2005) for PFt and BA44.

Since these cytoarchitectonic maps do not provide labels in the lateral occipital lobe, we used in this case the lateral occipital cortex in the Harvard-Oxford atlas (https://fsl.fMRIb.ox.ac.uk/fsl/fslwiki/Atlases), which provided a good approximation to the cluster yielded by the localizer ISC in the lateral occipital lobe, and did not include any other cluster of activity identified in the localizer ISC (See Fig. 2a). The Juelich maps and the lateral occipital cluster were transformed from the MNI into the native fMRI space using the same procedure used for the localizer maps. The final regions of interest were computed as the conjunction between the localizer and the Juelich maps. Additionally, the cortical depth maps estimated in each subject’s whole-brain MP2RAGE scan were taken into the native space using the same procedure.

Using these ROIs, we extracted the time courses to feed into the second ISC. The time courses were extracted from the fMRI data in native space after basic preprocessing (motion correction + trend removal and no spatial smoothing). For each region of interest in PFt, BA44 and the latOCC region, the encompassed voxels were binned according to their cortical depth. For each cortical depth bin, we concatenated the segments of the time course containing movie stimuli, transformed the intensity values in Z-scores and averaged the resulting values across voxels (as in the first ISC). This resulted in a time course containing the brain activity elicited by the movies for each cortical depth bin, each region of interest and each subject.

Given the nominal resolution of our data (0.8 mm), it is not possible to bin the cortical depth to selectively isolate the fMRI signal specific to each cortical layer. However for our experiment we did not aim to estimate the specific activity of each layer, but rather to estimate differences between intact and scrambled movies as a function of depth. For this reason, as recommended by Uludag and Havlicek (Uludag and Havlicek 2021), rather than using the acquisition resolution, we carried out the second ISC analysis after sampling the cortical depth in 6 depth bins. In supplementary materials we further verify that results look similar when using 8 and 10 bins. Correction for multiple comparisons (using false discovery rate - FDR) was adapted to the choice of number of bins. Inference on the difference between the ISC values for Intact and Scrambled in each cortical depth bin and for each region of interest was carried out using a paired t-test on the r->z transformed ISC values after verifying normality using the Shapiro-Wilk test. For illustration in figures, we then z->r backtransformed the values to provide the reader with the more familiar r values.

### ISFC

In addition to the ISC, we performed a depth-resolved inter-subject functional connectivity (ISFC) analysis. In this analysis, we compute the functional connectivity across each bin (bin 1 - close to the GM/WM interface - to bin 6 - close to the pial surface) and each region of interest (PF, BA44 and latOCC). We compute this functional connectivity with a leave-one-out ISFC approach. Precisely, this procedure consists in computing the correlation of the time course associated to *bin*_*j*_ in the region of interest *JU*_*i*_ for subject *s*, with the leave-one-out average time course associated to *bin*_*l*_ in the region of interest *JU*_*k*_ (averaged over all subjects except *s*). As a result, we have for each combination of *bin* and *JU* and for each subject *s* an estimate of the intersubject functional connectivity between *bin*_*j*_ in *JU*_*i*_ and *bin*_*l*_ in *JU*_*k*_ :

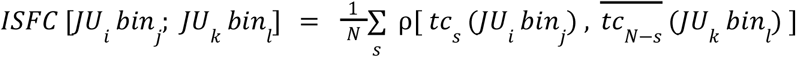

Where *tc*_*s*_ (*JU*_*i*_ *bin*_*j*_) is the time course associated *bin*_*j*_ in *JU*_*i*_ for subject *s* and *ρ* is a Pearson correlation.

We transformed such correlation values to Z scores (using Fisher r-to-Z transformation) and used them as input for a one sample t-test to assess whether each condition exhibits non-null functional connectivity across these regions. Additionally, we perform a paired sample t-test between the intact and scrambled condition, to determine whether the specific condition has an impact on the functional connectivity that we determine with ISFC.

Alongside the ISFC, we carry a more traditional PPI-like functional connectivity analysis (Friston et al. 1997). In this approach, we perform all the correlations in a within-subject fashion and subsequently compute statistics across subjects, i.e.,

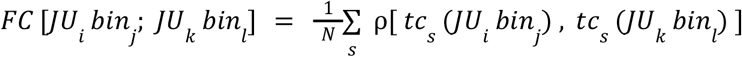

Where *tc*_*s*_ (*JC*_*i*_ *bin*_*j*_) is the time course associated *bin*_*j*_ in *JC*_*i*_ for subject *s* and *ρ* is a Pearson correlation. Inference on the difference between the ISFC values for Intact and Scrambled in each cortical depth bin and for each region of interest was carried out using a paired t-test.

## Results

### Premotor, parietal and lateral occipital cortices show significant ISC for intact or scrambled sequences

First, we aimed to macroscopically localize premotor, parietal and posterior temporal/lateral occipital patches of interest with activity that carries movie-related information in intact or scrambled movies. To do so, we transformed the preprocessed 4D fMRI data into the MNI space, smoothed the BOLD signal, and calculated the leave-one-out ISC for each voxel separately for intact and scrambled sequences. This revealed the expected network including the lateral premotor cortex and BA44, the primary somatosensory cortex and the inferior parietal lobule (including PFt) as well as lateral extrastriate occipital regions (latOCC) (Fig. 2). This network resembles that found using the same stimuli in conventional resolution 3T fMRI (Thomas et al. 2018). We then identified voxels containing significant ISC for the intact or scrambled movie in PFt, BA44 and latOCC as regions of interest, to then resolve the ISC as a function of cortical depth. By selecting them with a logical OR between the results of the ISC carried out either on intact or scrambled movies (i.e. taking all voxels surviving the threshold for ISC_intact_ >0, or for ISC_scrambled_ > 0), we avoid biasing the regions of interest towards intact or scrambled movies in further analyses (Kriegeskorte et al. 2009).

### PFt shows increased stimulus related information at depths corresponding to layers III,V and VI for intact > scrambled sequences

The second ISC analysis aimed at resolving the cortical depth of the fMRI BOLD signal synchronization in PFt for Intact vs Scrambled movies. Therefore, this analysis was carried out using the preprocessed but unsmoothed data in the native fMRI space of each participant. For each subject an average time course was calculated across all voxels within each depth-bin in PFt. Then, a leave-one-out ISC was carried out for each bin, i.e. the signal of each subject *i* in a depth-bin was correlated with the average signal of all other subjects in the same depth bin of the same ROI.

The results of this depth-resolved ISC for PFt are displayed in Figure 3a and show how the difference in stimulus locked signals between the observation of Intact and Scrambled movies depends on the cortical depth. As predicted (Fig. 1), during the observation of Intact movies, the synchronization of the activity in deep layers (at depths aligning with layers V/VI) of PFt significantly increased. We also observed significant increases of ISC at depths aligning with layer III, in line with the tracer studies in monkeys that suggest that feedback from BA44 to PFt also terminates in layer III. Draining vein effects might also contribute to this increase in layer III synchronization, and these effects cannot be disentangled with BOLD (Huber et al., 2017). For completeness, although we did not have specific hypotheses for ISC in BA44 and LatOcc, we also present the differential ISC as a function of depths for these regions in Supplementary Fig. S2. For BA44, the activity was significantly more synchronized in the superficial layers for the intact, and in the deepest layers for the scrambled movies (See Supplementary Fig S2). Similar results are obtained when the cortical depth was sampled in either 8 or 10 bins (See Supplementary Fig. S3). No significant changes were observed in our latOcc patch of interest, with the Bayesian analysis leaning towards the null hypothesis of no difference in all but one bin, suggesting that participants processed intact and scrambled movies similarly in the visual system. Examining the individual contrast values in PFt (Fig. 3b) shows that the individual participants showed relatively consistent effects in our sample. Finally, examining the actual ISC values in the two conditions (Fig. 3c and Supplementary Figure S2) confirms the general trend found in the literature, that ISC is highest in earlier sensory regions (latOcc peak average ISC around r = 0.3), lower in parietal cortices (PFt peak average ISC around r = 0.2), and lowest in frontal cortices (BA44 peak average ISC around r = 0.1).

**Fig 3.**
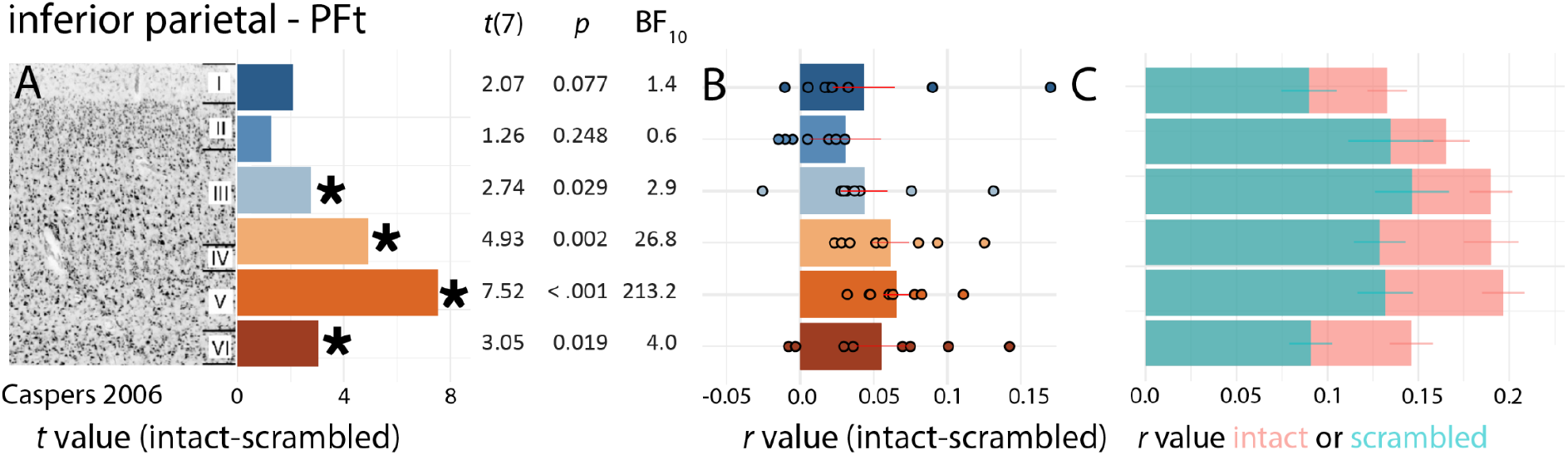
Depth-resolved ISC for the contrast Intact - Scrambled movies in PFt. (a) t-values of the contrast intact > scrambled in PFt as a function of depth bin, together with the two-tailed p and BF values all derived from a matched pair t-test of the r→z transformed leave-one-out ISC values per bin. Significant t-values after FDR correction (q(FDR) = 0.05) across the 6 bins are indicated with an asterix. Results are shown next to a histological reconstruction of the 6 cortical layers of PFt in postmortem brains from Caspers et al. (Caspers et al. 2006) for visual guidance only. The consistency between these results and those obtained with 8 or 10 bins is shown in Supplementary Figure S3. (b) The difference in leave-one-out r-value between intact and scrambled movies as a function of depth in PFt. Results were generated by Fisher r→z transformation before averaging and calculating s.e.m., and then back-transformed into r-values for illustration. For all bins in (a) we show the ISC value of each participant. Error bars are s.e.m. (c) r-value of the leave-one-out ISC as a function of bin for the intact (pink) and scrambled (green) condition separately.

### Functional connectivity is increased across PFt and BA44 for intact sequences

After we found evidence for increased ISC for Intact over Scrambled movies in PFt, we probed the interaction between PFt and either BA44 or latOCC in the two conditions. We hypothesized that during the observation of Intact movies, brain activity would be more enhanced in the feed-back stream from BA44 to PFt, terminating at depth aligning with layers III and/or V/VI (Gerbella et al. 2011). Within the visual system, deep-layers are particularly associated with predictive brain activity (Shipp 2007; Kok et al. 2016; de Lange et al. 2018; Aitken et al. 2020), but whether the same is true in the fronto-parietal system was unclear. In contrast, for the information transfer across latOCC and PFt, because the latOCC appears to represent information at the short temporal scale of acts, too fast for the the Nyquist limit of our acquisition, we do not expect measurable changes in ISFC. Had we had higher temporal resolution, we might have expected the scrambled condition in particular, to increase ISFC from latOCC towards the intermediate depth of PFt aligning with the granular layers.

Fig. 4 (left column) represents the results of the ISFC. In accordance with our hypothesis we found the information transfer, as assessed using ISFC, between PFt and BA44 to be increased for intact compared to scrambled conditions (Fig. 4a). Comparing the depth bins with significant ISFC changes with the anatomy of PFt and BA44 suggests that ISFC is most increased across layer III in PFt and BA44, although a significant increase was also found between depths corresponding to layer VI in PFt and layers I-II in BA44. In line with the idea that the additional exchange of information between PFt and latOCC conveying prediction errors in the scrambled condition would occur at a time-scale too fast for our acquisition to capture, we did not find a significant contrast in ISFC across conditions between these regions. Examining the ISFC values in the individual conditions for PFt x BA44 (Fig. 4 left middle and bottom row) confirms that although some significant ISFC occurs in all conditions, the intact condition stands out with more consistent ISFC in the intact condition.

**Fig. 4.**
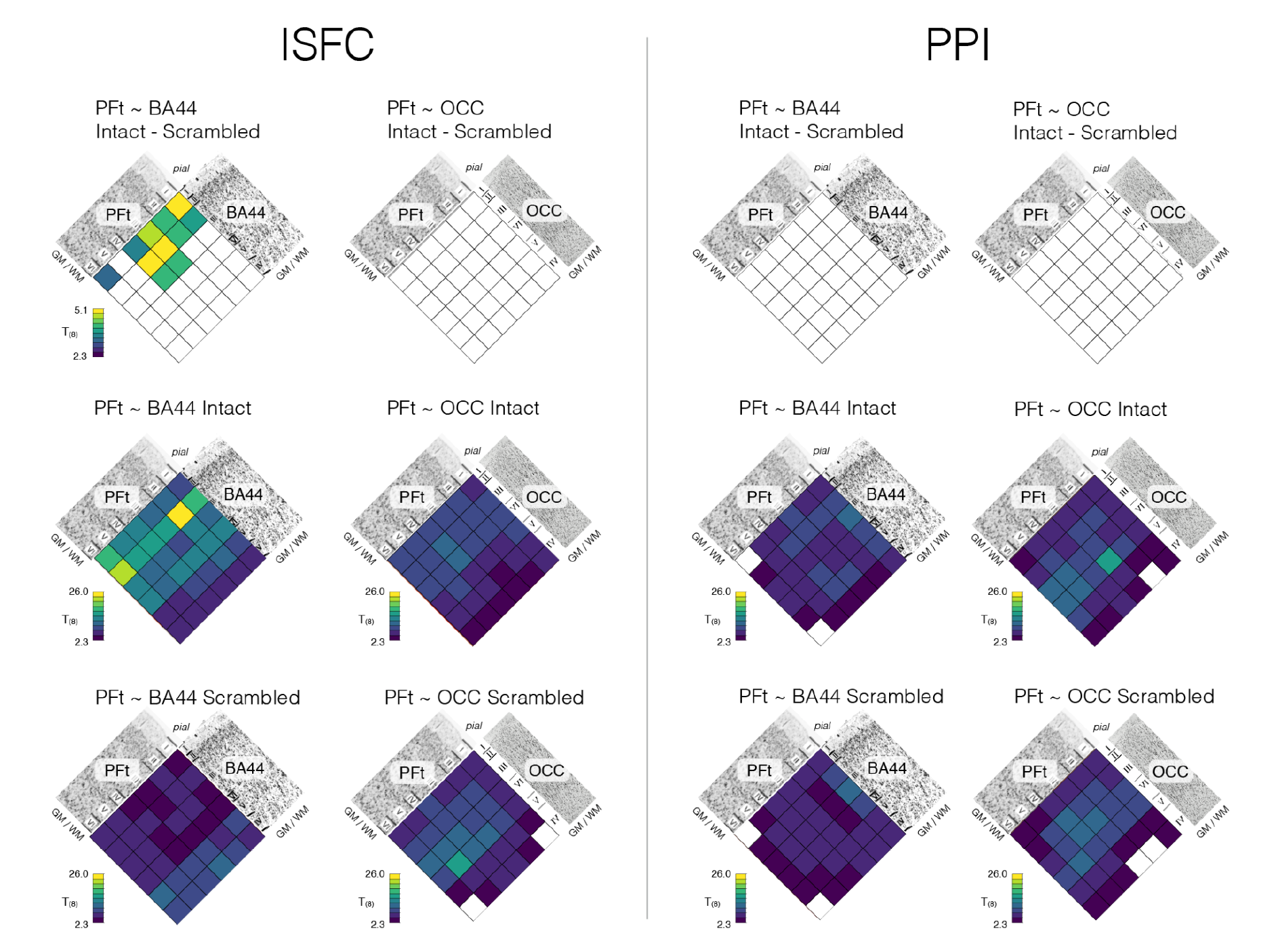
Results of the ISFC. Left column: Significant ISFC differences for the Intact - Scrambled contrast (top row), as well as for the Intact>0 (middle) and Scrambled>0 (bottom) contrast separately. **Right column:** Same but using a traditional PPI analysis using the same dataset. Note: ISFC calculates the correlation between time-courses in region A of each of the N participants and region B averaged across the other participants, and then contrasts these N values across conditions using a paired t-test. PPI calculates the correlation between regions A and B in the same participant for each of the N participants, and then contrasts these N values across conditions. The same color scale was used throughout to permit the comparison of t-values across the two methods. The intersubject variability of PFt ∼ BA44 Intact - Scrambled ISFC is displayed in Fig. S4.

To illustrate the advantage of ISFC over traditional within-subject functional connectivity, we used the same data, but calculated the correlation between the time course of one area and that of another within each participant, rather than across a participant and the average of the others (Fig. 4 right column). Contrasting this within subject correlation across our two conditions is then equivalent to a traditional PPI (Psycho-Physiological Interaction(Friston et al. 1997)), but could not reveal a specific change in our sample. The advantages of ISFC over a traditional PPI are in line with previous studies and is thought to be due to the fact that calculating correlations across participants rather than within participants reduces the impact of common noise across voxels(Simony et al. 2016; Nastase et al. 2019).

## Discussion

Our aim was to shed further light onto the functional architecture of action observation by providing depth-resolved fMRI data that could help evaluate one of the core notions of predictive coding: that parietal nodes receive increased feed-back from premotor region BA44 when actions organize into predictable sequences. Specifically, we wanted to test whether sequence level information - isolated through the contrast intact-scrambled - (1) can be found in layers of PFt known to receive feedback connections from BA44, namely layers V/VI and III (Gerbella et al. 2011), and (2) is shared with layers of BA44 that could be sending feedback to PFt (layers II/III and V/VI). The analytical methods of ISC and ISFC were used to tackle the first and the second question, respectively.

As in our previous 3T MRI study (Thomas et al. 2018), at the macroscopic level we found that premotor, parietal and lateral occipital clusters of voxels encoded information about the observed motor acts, as measured by significant intersubject correlation while viewing the intact *or* scrambled movies. This was true whether we used non-parametric methods advised in the literature(Chen et al. 2016), or parametric methods we have recently shown to be more sensitive while at the same time controlling for Type-I errors(De Angelis et al. 2021). We then focused on PFt because predictive coding models and traditional feed-forward models of action observation make predictions that differ regarding the recruitment of deep layers in PFt: only such models predict that feed-back information should increase in intact over scrambled sequences, and hence that the contrast intact > scrambled should reveal increased ISC in layer III or V/VI of PFt. To focus on voxels in PFt that represent the movies, we thus transformed the coordinates of the MNI voxels with significant information in at least one of the conditions at the group level (intact or scrambled) into the participants’ native, unsmoothed space to define our PFt patch of interest. This patch was not defined based on a particular relationship between intact and scrambled ISC, and is thus not biased towards either condition(Kriegeskorte et al. 2009). In native space, using unsmoothed data, we then calculated the leave-one-out ISC between a participant’s average activity in that bin and the average of the other participants in the same bin, and compared ISC in the intact and scrambled condition. As in our previous 3T study(Thomas et al. 2018), we found that PFt showed higher ISC while participants viewed intact than scrambled sequences. Importantly, for the first time we resolved the depth-profile of such sequence-level information. In accordance with predictive coding and the notion that predictive top-down information should be present in deeper layers (Kok et al. 2016; de Lange et al. 2018; Aitken et al. 2020), we found the significant difference with ISC intact > ISC scrambled to peak in the deeper parts of PFt, at a depth that aligns with layer V in postmortem histological examinations of the brains of other individual(Caspers et al. 2006). In line with anatomical tracing studies in monkeys, suggesting that feedback connections from BA44 to PFt should terminate in layer III in addition to layers V/VI, this increased ISC was significant also at depth compatible with the location of layer III. Although we had no particular hypotheses for the depth profile of latOcc and BA44, as in our previous 3T study, we found that latOCC does not show significant differences in ISC between intact and scrambled sequences, irrespective of cortical depth, while BA44 did show significantly more ISC for intact vs scrambled conditions in superficial layers. See Supplementary Note 1 for further details. Importantly, that latOCC shows high ISC values of similar strength for the intact and scrambled condition shows that participants engaged with the material under both conditions, and that differences in regions such as PFt or BA44 are unlikely to be due to reduced information processing in visual cortices.

By combining our comparatively higher spatial resolution with ISFC - to avoid the issue of shared noise across brain regions(Simony et al. 2016; Nastase et al. 2019) - we could also verify the prediction that the sequence level information at PFt depths aligning with layers III, V and VI is shared with BA44 but not with LatOcc. Because this increased ISFC is calculated across different subjects, it indicates that it is signal relating to the intactness of the sequence in the stimuli that synchronizes across BA44 and PFt (Simony et al. 2016; Nastase et al. 2019). The depth of increased ISFC across PFt and BA44 suggests that feedback information was most strongly increased across supragranular layers between PFt and BA44. This is compatible with findings of tracing studies in macaque monkeys that show connections from BA44 terminating in layers III (Gerbella et al. 2011), and data showing that feedback connections can originate from layers II/III (Shipp 2007). However, a smaller increase was also observed between superficial layers of BA44 and the deepest depth of PFt. In contrast, in the visual system proper, predictive processes were mainly accompanied by an increased recruitment of the deeper layers of the cortical visual system (Kok et al. 2016; de Lange et al. 2018; Aitken et al. 2020). This difference is perhaps not surprising given that the prototypical pattern of feed-forward vs feed-back connectivity originally described by Felleman and Van Essen (1991) is clearest in early sensory brain regions, but becomes increasingly columnar or idiosyncratic in association brain regions and the motor system (Finn et al. 2020). Performing such studies in association regions, as we have done here, may thus be critical to shed light on how easily results from the much studies visual system apply to other cortical loops.

Although it is difficult to equate BOLD signal at a certain cortical depth with activity in a particular cortical layer, it should be noted that the depth bins aligning with layer III in PFt, in which we measure increased ISC and ISFC for intact sequences, inevitably also partially aligns with the depth of the thin layer IV in PFt. This makes it impossible to verify our prediction that layer IV, thought to receive feed-forward information from visual regions, should not show increased ISC or ISFC for the Intact sequences. Even increasing the number of bins in our depth analysis cannot overcome this issue for two reasons: first, because capillaries flow through layer IV to reach layers V, VI so that any increase in ISC/ISFC in neurons in layers V or VI will also increase the ISC/ISFC in the BOLD signal in layer IV (Stephan et al. 2019); second, because resampling data in an increasing number of depth-bins after acquisition at 0.8mm voxel size will lead to vertical blurring.

Our study has a number of additional limitations. First, as often at high-field fMRI, we used fewer participants scanned twice rather than more participants (Baker et al. 2020; Cai et al. 2021) due to the difficulty of finding participants that can limit head movement to ensure stability at 0.8mm resolution. This may limit the degree to which we can extrapolate our findings to the general population, although the similarity of our results with those of a larger group acquired at 3T at the macroscopic level (Thomas et al. 2018), and the limited spread of our participant’s r-values illustrates that the effects we report were quite robustly observed across participants. Second, to maintain our processing pipeline as close as possible to the actual data, we did not attempt to remove the bleeding of activation from deeper to more superficial layers due to the organization of the cortical vascularization (Markuerkiaga et al. 2016; Stephan et al. 2019; Uludag and Havlicek 2021). Future studies using more elaborate models of how blood flows through the depth of the cortex may provide a clearer picture on whether signal changes in the intermediate levels aligning with layer IV across PFt and BA44 do or do not lead to local changes in ISFC. Third, the comparatively slow acquisition rate of our fMRI signal makes it impossible for us to tune accurately in the prediction error signals that we expect to occur at each camera change in our datasets. We are currently analysing data acquired using ECoG in epileptic patients to try to shed light on this faster temporal scale.

Finally, although we use depth resolved fMRI to quantify the level of feed-back input to PFt across conditions in ways that have become relatively frequent in the 7T literature, attributing activity at a particular depth to feed-back vs feed-forward information flow hinges (a) on the separability of these information flows along the cortical depth and (b) on our knowledge of where these inputs terminate. While in primary sensory regions such as V1 both these conditions appear to be met, with feed-forward signals peaking in Layer IV while feedback signals alter activity elsewhere (Self et al. 2019), the connectivity pattern between parietal and premotor regions in monkeys are, as we mentioned, more complex, with some frontal input to parietal regions terminating throughout all layers in a lateral connectivity pattern while others are multilayered, avoiding layer VI, albeit with a pattern that differs from the prototypical pattern described in early visual regions (Felleman and Van Essen 1991; Rozzi et al. 2006; Shipp 2007; Gerbella et al. 2010, 2011). While studies like ours have thus been argued to be necessary to start exploring whether the prevalence of predictive feed-back signals in deep layers observed in the visual system applies elsewhere (Finn et al. 2020), there is also urgent need to perform the kind of laminar recordings that have benchmarked the foundations of this approach in the visual system(Self et al. 2019) across posterior parietal and premotor regions.

Despite these limitations, we hope that this study will serve as a proof of concept that depth-resolved analyses can be performed during naturalistic viewing in higher cognitive brain regions using intersubject correlation-based approaches, and that we may contribute to what Finn and colleagues called for: “layer fMRI is now at a point where we can expand from tightly controlled experiments in sensory cortex with clear hypotheses—which were necessary to show feasibility of the technique—to more exploratory, data-driven investigations of functional dynamics both across the cortical hierarchy as well as within higher-order regions themselves. […] Data acquired during naturalistic stimulation—e.g., movie watching—lend itself to both connectivity and activation analyses” (Finn et al. 2020). Ultimately such studies may literally contribute to a deeper understanding of social cognition.

## Author contributions

Following the Contributor Roles Taxonomy (CRediT) available at https://casrai.org/credit/. The relative amount of markers highlights the degree of contribution within each role.

| Author | RB | LC | LdA | WvdZ | PLB | VG | CK |
| --- | --- | --- | --- | --- | --- | --- | --- |
| **Conceptualization** | ** |  |  |  |  | *** | *** |
| **Data curation** | * | *** |  |  |  |  |  |
| **Formal analysis** | * | *** | ** |  |  |  |  |
| **Funding acquisition** | *** |  |  |  |  | *** | *** |
| **Investigation** | *** |  |  | ** |  |  |  |
| **Methodology** | *** | *** | *** | ** | ** | *** | *** |
| **Project Administration** | ** | ** |  |  |  | *** | *** |
| **Resources** | *** |  |  |  |  | *** | *** |
| **Software** |  | *** | ** |  | ** |  |  |
| **Supervision** |  | * |  |  |  | *** | *** |
| **Validation** |  | *** | *** |  |  |  |  |
| **Visualization** |  | *** |  |  |  | * | * |
| **Writing original draft** | * | *** | ** |  |  | *** | *** |
| **Writing review and editing** | * | * | * | * | * | * | * |

## Acknowledgements

We thank Giuseppe Luppino for guiding our hypotheses about the layers of PFt that should receive BA44 input based on the tracing studies in monkeys from his lab, Teresa de Sanctis for help with preparing the stimuli, the members of the Spinoza centre where the scanning was conducted for providing technical support, Robert Turner for several helpful discussions on depth specific 7T fMRI.

## Supplementary Online Materials

Supplementary Figure S1 : Field of view of the fMRI acquisitions

**Figure S1:**
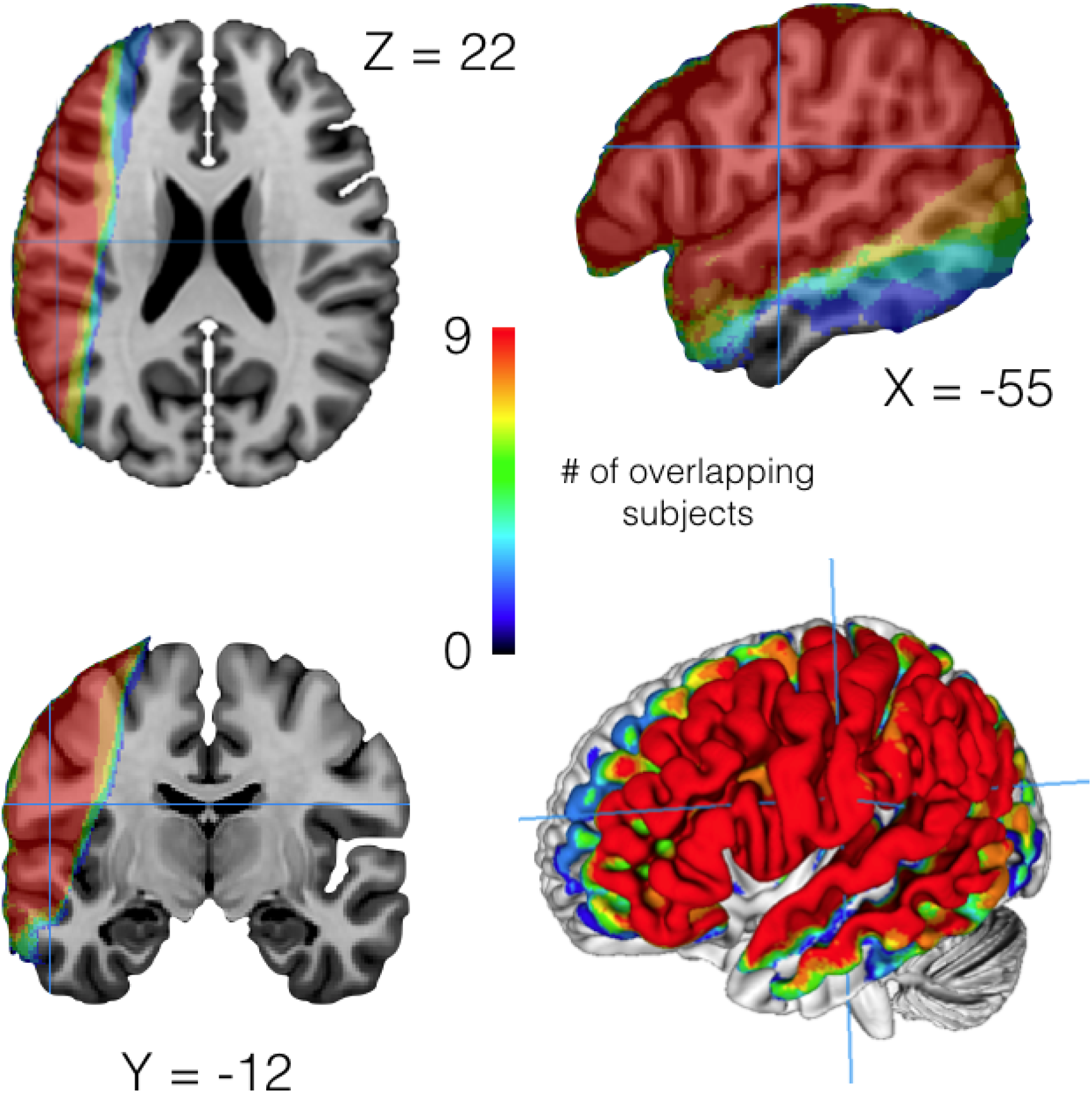
Coverage of the left hemisphere. We centered our planning of the 7T acquisition on the parietal area PFt. Because the planning was done subject by subject, not the exact same coverage in terms of MNI coordinates was achieved in all participants. Here we therefore show the overlap in coverage after transforming the field of view into MNI space. As can be seen, we achieved consistent coverage in inferior parietal and frontal regions as well as in anterior lateral occipital cortex.

### Supplementary Note 1 and Figure S2: ISC as a function of depth in LatOcc and BA44

Although we had no particular hypotheses for the depth profile of latOcc and BA44, as in our previous 3T study (Thomas et al. 2018), we found that latOCC does not show significant differences in ISC between intact and scrambled sequences, irrespective of cortical depth (Supplementary Figure S2a), with Bayes factors leaning in favour of the null hypothesis in most depth bins. This is in line with other studies showing that more anterior regions integrate information over longer periods than early sensory regions, be they visual or auditory (Lerner et al. 2011; Honey et al. 2012). In BA44 we found a more complex pattern of differences between intact and scrambled sequences. The two most superficial depth bins showed ISC that was higher in intact than scrambled conditions, while the deepest depth bin showed the opposite effect. While finding intact>scrambled in BA44 was expected, that this difference was most marked at superficial depth was not: based on previous findings in visual brain regions, one might have expected sequence-level information to be strongest in the depth of BA44 (Kok et al. 2016; de Lange et al. 2018; Aitken et al. 2020). Furthermore, that the deepest parts of BA44 would show more ISC for scrambled sequences was entirely unexpected and we feel uncertain about how to interpret this finding. Some participants reported that viewing the scrambled sequences created an oddly dissonant experience of cognitively understanding what action was being performed at a more abstract level despite the perceptual frustration of a visual sequence that constantly violated expectations in the timing of the acts. In a way, the sum of the acts made sense at an abstract level, but their sequence did not. Perhaps, one could thus speculate that this hybrid pattern in BA44, with relatively increased ISC in superficial voxels for intact and in deep voxels for scrambled sequences could represent a switch in the dominant processes, with the more superficial voxels processing sequence level information embedded in the temporal stream of the visual input (which is more reliable in the intact sequence) and deeper voxels processing top-down input from cognitive processing that is more detached from the temporal stream of the visual input, and becomes more critical when the visual input itself does not make sense (in the scrambled condition). However, such an interpretation is very tentative indeed.

It is also worth noting that the increase of ISC in the top most superficial bins aligns well with the fact that ISFC between BA44 and PFt is also increased in these two most superficial layers of BA44.

**Figure S2:**
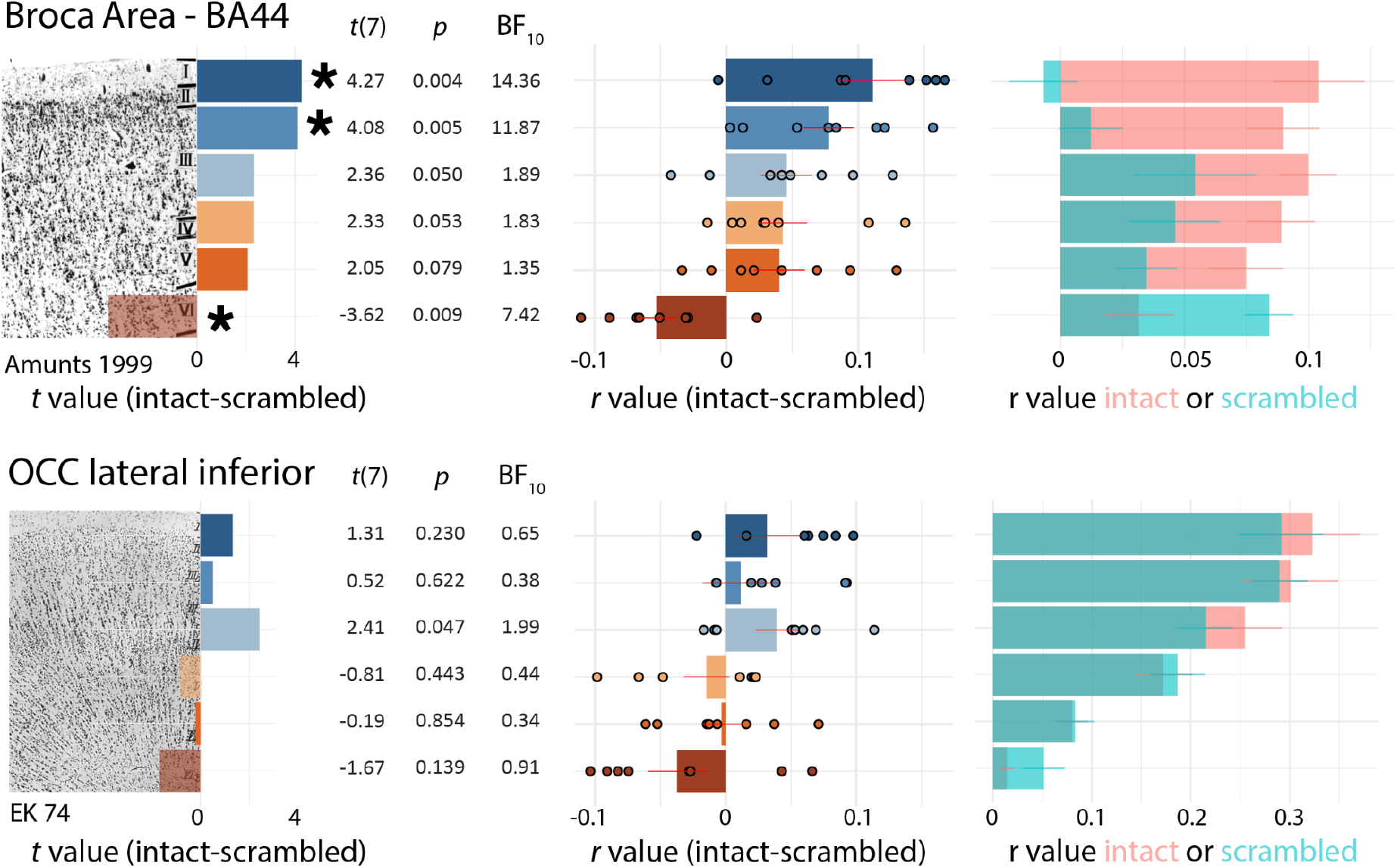
ISC as a function of depth in LatOcc and BA44. All conventions as in Figure 3, but for Lateral Occipital cortex, using a postmortem analysis from peristrate area OA also known as area 74 of Economo & Koskinas (EK74, von Economo 1927, as reprinted in 2009, p. 105) within LatOcc; or for BA44 using a postmortem analysis from Amunts et al.(1999). *t*(7), *p* and BF_10_ values all stem from a two-tailed, matched pair t-test on the r->z transformed ISC values for intact vs scrambled. P values are uncorrected for multiple comparisons but asterisks denote bins at which the *p* value after fdr correction is below 0.05 (*q*<0.05).

Supplementary Figure S3 : Consistency of ISC results across binning

**Fig. S3.**
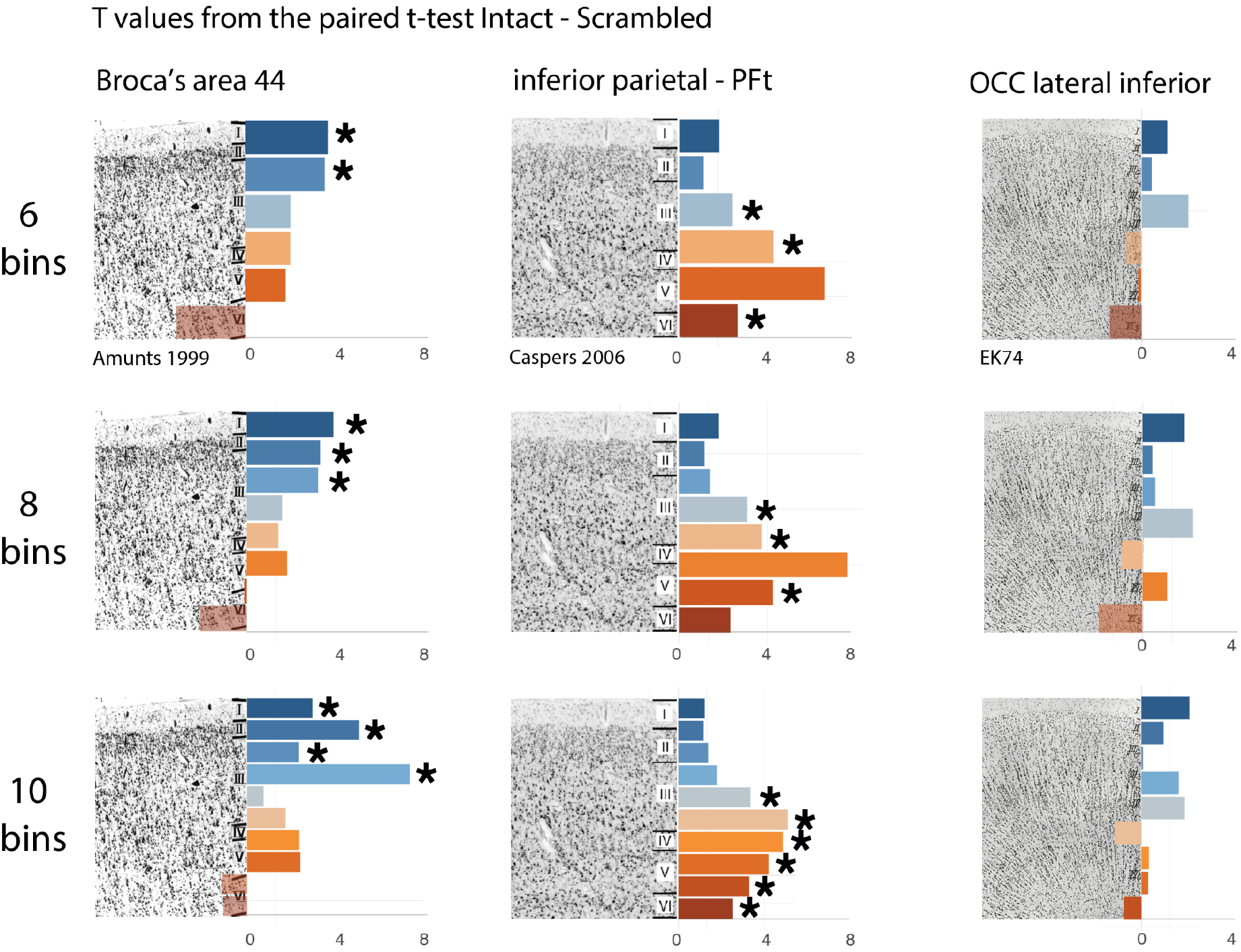
Consistency across binning choices. Bars represent the t value from comparing the ISC estimates in either condition using a paired t-test. Significance was determined using FDR correction (q(FDR) = 0.05 - relative to the number of bins) across all bins within each region, and is indicated by the asterisk.

Supplementary Figure S4 : Intersubject variability in ISFC

**Fig. S4.**
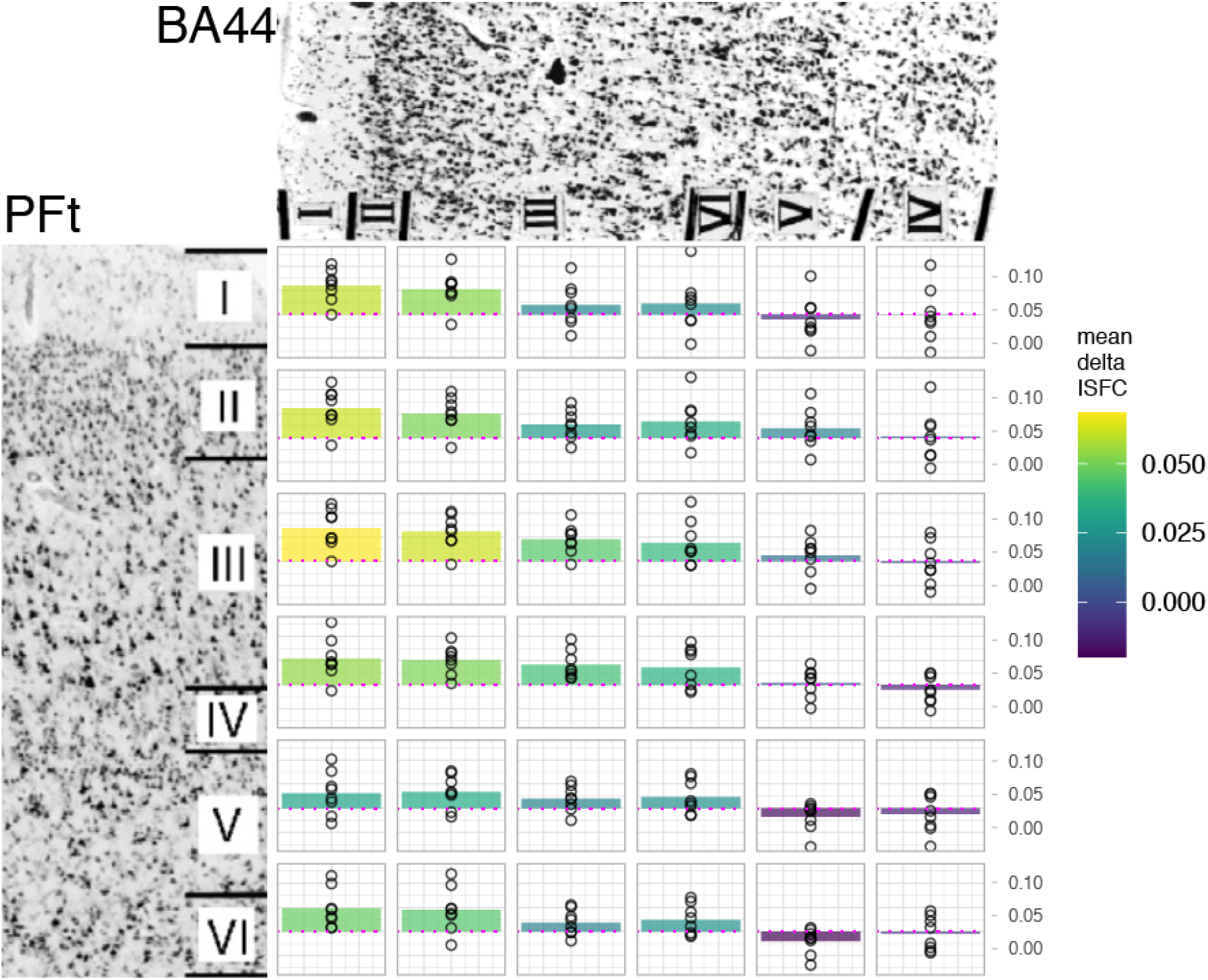
Intersubject variability for significant values of the PFt ∼ BA44 ISFC in the Intact - Scrambled contrast. Mean Delta ISFC denotes the mean difference between ISFC *r* values in the Intact and the Scrambled conditions. This value is used to color-fill the barplots. Each circle represents one participant’s ISC *r*-value value. The dotted magenta line highlights the zero.

### Supplementary Materials 5 : Quality control images for preprocessing steps: skull stripping, registration, transformation of localizer/atlas following registration, layering

This folder of the github repo (https://github.com/ldeangelisphys/layerfMRI/tree/main/QC_images) contains quality control images documenting the outcome of the following procedures:

| Procedure | Example QC image |
| --- | --- |
| summary skullstrip: outcome of the skull-stripping for the partial FOV fMRI and low-res anatomical, as well as of the high-res anatomical image. | https://github.com/ldeangelisphys/layerfMRI/blob/main/QC_images/summary_skullstrip/sub_02_skullstrip.pdf |
| summary registration and summary inv_registration: registration between different spaces: fMRI native (4 partial FOV sessions per subject), low-res anatomical, high-res anatomical, MNI | https://github.com/ldeangelisphys/layerfMRI/blob/main/QC_images/summary_registration/sub_02_registration.pdf<br>https://github.com/ldeangelisphys/layerfMRI/blob/main/QC_images/summary_inv_registration/sub_02_inv_registration.pdf |
| summary layering: high-res anatomical skull strip, cortical segmentation and cortical depth estimation | https://github.com/ldeangelisphys/layerfMRI/blob/main/QC_images/summary_layering/sub_02_layering.pdf |
| summary trans2native: transformation of the cortical depth maps into the native fMRI space | https://github.com/ldeangelisphys/layerfMRI/blob/main/QC_images/summary_trans2native/sub_02_native_space.pdf |

